# Transmembrane 163 (TMEM163) protein interacts with specific mammalian SLC30 zinc efflux transporter family members

**DOI:** 10.1101/2022.07.07.499096

**Authors:** Adrian Escobar, Daniel J. Styrpejko, Saima Ali, Math P. Cuajungco

## Abstract

Recently, we reported that TMEM163 is a zinc efflux transporter that likely belongs to the mammalian solute carrier 30 (Slc30/ZnT) subfamily of the cation diffusion facilitator (CDF) protein superfamily. We hypothesized that human TMEM163 forms functional heterodimers with ZNT proteins based on their subcellular localization overlapping with TMEM163 and previous reports that certain ZNT monomers interact with each other. In this study, we heterologously expressed individual constructs with a unique peptide tag containing TMEM163, ZNT1, ZNT2, ZNT3, and ZNT4 (negative control) or co-expressed TMEM163 with each ZNT in HEK-293 cells for co-immunoprecipitation (co-IP) experiments. We also co-expressed TMEM163 with two different peptide tags as a positive co-IP control. Western blot analyses revealed that TMEM163 dimerizes with itself but that it also heterodimerizes with ZNT1, ZNT2, ZNT3, and ZNT4 proteins. Native co-IP using mouse tissues confirmed the interactions while confocal microscopy revealed that TMEM163 and ZNT proteins partially co-localize in cells, suggesting that they exist as homodimers and heterodimers in their respective subcellular sites. Functional zinc flux assays using Fluozin-3 and Newport Green dyes show that cells expressing TMEM163 homodimers extruded zinc slightly less efficiently than cells expressing TMEM163/ZNT heterodimers. Cell surface biotinylation revealed a subtle change in the plasma membrane localization of TMEM163 upon co-expression with certain ZNT proteins, which possibly explains why zinc efflux is marginally different for TMEM163 homodimers than TMEM163/ZNT heterodimers. Overall, our results show that the interaction between TMEM163 and distinct ZNT proteins is functionally relevant and that their heterodimerization may serve to influence their zinc efflux activity within specific tissues or cell types.

**Research Highlights:**

- TMEM163 protein heterodimerizes with mammalian ZNT1, ZNT2, ZNT3 and ZNT4 zinc efflux transporters.
- TMEM163 and ZNT proteins partially co-localize in their respective plasma membrane or subcellular compartments, suggesting distinct cellular roles as homodimers and heterodimers.
- The zinc efflux activity of TMEM163 or ZNT protein homodimers did not markedly differ from their TMEM163/ZNT heterodimer counterparts.
- Functional TMEM163/ZNT heterodimers give further credence to the role of TMEM163 as a *bona fide* member of the SLC30 protein family

## 1. INTRODUCTION

Rodent Transmembrane 163 (Tmem163) protein was initially coined as synaptic vesicle 31 (Sv31), based on its identification from rat brain synaptosomes using proteomics and calculated molecular weight (Burre *et al.*, 2007). Human TMEM163 and mouse Tmem163 transcripts are detected in many tissues, notably in the brain, lung, pancreas, kidney, ovary, and testis (Cuajungco *et al.*, 2014; Sanchez *et al.*, 2019; Chakraborty *et al.*, 2020). Heterologous expression of TMEM163 in cultured HEK-293 cells exhibits localization within plasma membrane (PM), lysosomes, and vesicular compartments (Cuajungco *et al.*, 2014; Sanchez *et al.*, 2019; Chakraborty *et al.*, 2020). Similarly, immunocytochemistry and subcellular fractionation studies of cultured PC-12 cells stably expressing Tmem163 show that it is detected in the PM, lysosomes, early endosomes, and synaptic vesicles (Barth *et al.*, 2011). It was also reported that Tmem163 localizes within platelet dense granules (DG) and insulin granules, suggesting an important role in DG biogenesis (Yuan *et al.*, 2021) and insulin packaging or secretion (Chakraborty *et al.*, 2020). Rodent Tmem163 has been shown to bind and transport zinc as a dimer in a proton-dependent manner (Burre *et al.*, 2007; Barth *et al.*, 2011; Waberer *et al.*, 2017) and we recently reported that human TMEM163 is a zinc efflux transporter (Sanchez *et al.*, 2019). Based on its function and subcellular localization, we proposed that TMEM163 be classified as a member of the mammalian ZnT (Slc30, solute carrier 30) subfamily of the cation diffusion facilitator (CDF) protein superfamily (Styrpejko and Cuajungco, 2021). The human ZNT (SLC30) family consists ten highly conserved, proton-dependent zinc transport proteins, namely, ZNT1 to ZNT10 (Eide, 2006; Colvin *et al.*, 2010; Kambe *et al.*, 2014). The ZNT proteins efflux zinc from the cytoplasm into the extracellular space and into the lumen of membrane-bound compartments. ZNTs have predicted six transmembrane domains (TMD), but the structure of one of its members, ZNT8, was recently solved using cryo-electron microscopy, confirming six TMD with intracellular N- and C-termini regions (Xue *et al.*, 2020).

Our interests in understanding the pathology of Mucolipidosis type IV (MLIV) disease, which is caused by the loss of Mucolipin-1 (TRPML1) protein function (Bargal *et al.*, 2000; Bassi *et al.*, 2000; Sun *et al.*, 2000), led us to discover that zinc dyshomeostasis is present in MLIV patient cells, as well as cell culture and mouse models of MLIV disease (Eichelsdoerfer *et al.*, 2010). We subsequently identified human TMEM163 protein as an interaction partner of the TRPML1 ion channel (Cuajungco *et al.*, 2014). Nevertheless, it is possible that the abnormal zinc metabolism in MLIV disease is possibly due to disrupted TMEM163 and TRPML1 interaction since TMEM163 is a zinc efflux transporter (Sanchez *et al.*, 2019) that shuttles zinc into lumenal compartments while TRPML1 is permeable to zinc (Dong *et al.*, 2008) that may serve as a lysosomal zinc release channel (Cuajungco and Kiselyov, 2017) especially in neurons (Minckley *et al.*, 2019). The loss of TMEM163 and TRPML1 interaction could explain intracellular zinc accumulation in MLIV cells, particularly that RNA interference (RNAi)-induced knockdown (KD) of TMEM163 and TRPML1 in HEK-293 cells shows cytoplasmic zinc elevation, while over-expression of both proteins exhibits an opposite effect (Cuajungco *et al.*, 2014). Further, KD or knockout (KO) of mouse Tmem163 in MIN6 cells and platelets, respectively, also produce intracellular zinc accumulation following zinc exposure (Chakraborty *et al.*, 2020; Yuan *et al.*, 2021). It is interesting to note that KD of TRPML1 in cultured HeLa cells also results in anomalous lysosomal zinc accumulation when these cells are exposed to zinc (Kukic *et al.*, 2013). This phenomenon is mediated by the SLC30A4 (ZNT4) zinc efflux protein (Kukic *et al.*, 2013). Taken together, ZNT4 and TMEM163 appear to influence zinc transport into the lysosomes (Kukic *et al.*, 2013), and the roles of each protein could explain the cellular zinc imbalance linked to TRPML1 dysfunction (Cuajungco and Kiselyov, 2017). We hypothesized that TMEM163 and ZNT proteins like ZNT4 form heterodimers based on several fronts. First, TMEM163 proteins localize where certain ZNT protein are also detected (Kukic *et al.*, 2013; Cuajungco *et al.*, 2014). Second, TMEM163 shares an evolutionary relationship with ZNT proteins being all members of the CDF zinc efflux protein family (Sanchez *et al.*, 2019; Styrpejko and Cuajungco, 2021). Finally, although most CDF proteins exist as homodimers, several ZNT monomers heterodimerize with each other. For example, ZNT5 and ZNT6 proteins are only functional as heterodimers (Suzuki *et al.*, 2005; Fukunaka *et al.*, 2009). Other heterodimeric combinations that were reported include ZNT3/ZNT4, ZNT3/ZNT10 and ZNT2/ZNT10 proteins (Salazar *et al.*, 2009; Patrushev *et al.*, 2012; Zhao *et al.*, 2016). Similarly, ZNT1, ZNT2, ZNT3 and ZNT4 also form heterodimers with each other (Lasry *et al.*, 2014; Golan *et al.*, 2015). Interestingly, the subcellular localization of ZNT1 homodimer, which is typically observed in the PM, changes mostly into vesicular or compartmental sites upon heterodimerization with ZNT3 and ZNT4, but only partially as ZNT1/ZNT2 heterodimers (Golan *et al.*, 2015). The respective heterodimers were reported to be functional using a Zinquin fluorescence assay that detects changes in cytoplasmic zinc levels (Golan *et al.*, 2015). In the current paper, we report that human TMEM163 protein heterodimerizes with ZNT1, ZNT2, ZNT3, and ZNT4.

## 2. MATERIALS AND METHODS

### 2.1 Materials

We purchased co-immunoprecipitation kits (Pierce), control agarose resin (Pierce), T-Per tissue lysis buffer (Thermo Scientific), n-Dodecyl-β-D-maltoside (DDM) (Thermo Scientific, Waltham, MA), poly-D-lysine (PDL, Sigma-Aldrich), Sulfo-NHS-LC-LC-Biotin (Pierce) and Neutravidin agarose beads (Pierce) from Fisher Scientific (Waltham, MA). Culture media (Dulbecco’s Modified Eagle’s Media [DMEM] with 4.50 g/L glucose, 0.58 g/L L-glutamine, 0.11 g/L sodium pyruvate; Corning) and supplements (fetal bovine serum [FBS]; 100 mM sodium pyruvate; 100X Glutamax™; Gibco) were also obtained from Fisher Scientific. Primary antibodies were purchased from the following sources: anti-HA rabbit polyclonal antibody (pAb; #NB600-345, Novus Biologicals, Englewood, CO; #F3165, Millipore-Sigma, St. Louis, MO), anti-DDK (M2 Flag) mouse monoclonal antibody (mAb; NBP2-43570, Novus Biologicals; #H6908, Millipore-Sigma), anti-TMEM163 rabbit pAb (#NBP1-06608, Novus Biologicals), anti-Znt1 rabbit pAb (#PA5-77768, Thermo Scientific), anti-Znt2 rabbit pAb (#PA5-99761, Thermo Scientific), anti-Znt3 mouse mAb (#TA501498, Origene Technologies, Rockville, MD), and anti-Znt4 rabbit pAb (#PA5-80028, Thermo Scientific,). Secondary anti-mouse and anti-rabbit antibodies conjugated with infrared (IR)-Dye™ 800CW were purchased from LI-COR Biosciences (Lincoln, NE), while those antibodies conjugated with Alexa Fluor-488 and Alexa Fluor-568, respectively, were purchased from Fisher Scientific. Fluorescent zinc dyes, Fluozin-3 tetra-potassium salt (membrane impermeant, #F24194, Thermo Scientific) and Newport Green DCF acetate (#N7791, Thermo Scientific) were purchased from Fisher Scientific.

### 2.2 Cell culture

Human embryonic kidney (HEK)-293T and HeLa cells were purchased from American Type Culture Collection (ATCC; Manassas, VA). The cells were cultured without antibiotics in DMEM supplemented with 10% fetal bovine serum (FBS; Thermo Scientific, Waltham, MA). The cells were maintained in a standard humidified 37°C incubator supplied with 5% CO_2_. For functional zinc flux assays, we added 1 mM sodium pyruvate and 1X Glutamax™ as additional supplements in the media or buffer to improve cell viability during the experimental trials.

### 2.3 Recombinant DNA constructs

We used the In-Fusion™ (IF) HD cloning system online primer design website (Takara Bio USA, Mountain View, CA) to create primer sets (**Table S1**) for cloning and/or subcloning of open-reading frame (ORF) of TMEM163 (purchased from Origene Technologies), SLC30A1/ZNT1 purchased from Origene Technologies), SLC30A2/ZNT2 (variant 1, purchased from Origene Technologies; variant 2, a kind gift from Dr. Shannon Kelleher, University of Massachusetts, Amherst), SLC30A3/ZNT3 (a kind gift from Dr. Robert Palmiter, University of Washington, Seattle), and SLC30A4/ZNT4 (purchased from Origene Technologies). We used IF cloning for the pBI dual expression vector containing a minimal CMV promoter (Takara Bio USA) (**Fig. S1**). Specifically, we created two sets of pBI clones – one group consisting of pBI constructs with single open reading frame (ORF) of TMEM163, ZNT1, ZNT2, ZNT3, and ZNT4; and another group that contains two cDNAs consisting of TMEM163 and one of each ZNT. For single expression of pBI constructs, the TMEM163 coding sequence was cloned at the second multiple cloning site (MCS2) leaving MCS1 empty, while the ZNT proteins were placed at the other site (MCS1) leaving MCS2 empty. For dual expression of the pBI constructs, each ZNT ORF was cloned in the MCS1 of the pBI vector containing TMEM163 in its MCS2. The use of pBI dual expression vector minimizes confounding variables arising from low transfection efficiency or biased expression for one of the two plasmids transfected, and potential cytotoxic effects of two-plasmid over-expression with CMV promoters. For protein-protein interaction experiments using the pCMV6 vector with HA or Myc-DDK peptide tag at the 3’-end (Origene Technologies), we used the same IF cloning system to subclone cDNAs of our genes-of-interests (GOIs) (**Table S1**). If the IF cloning approach failed, we cloned the GOIs into the pCMV6 vector using restriction enzymes, *Sgf I* at the 5’ end and *Mlu I* at the 3’ end, since we designed the IF cloning primers containing these restriction sites. For TMEM163 and ZNT4, we subcloned their ORFs into pCMV6 vector with HA or DDK tag directly from a commercially purchased pCMV6-GFP and pCMV6-HA expression clones (Origene Technologies), respectively, using *Sgf I* and *Mlu I* restriction enzymes. The sequence integrity of all expression constructs used in the study was verified by Sanger sequencing (Retrogen, San Diego, CA) and analyzed using Lasergene SeqMan version 17 (DNAStar, Madison, WI).

### 2.4 Co-immunoprecipitation (Co-IP) and Western blot (WB)

HEK-293T cells were seeded at 1.5 × 10^6^ cells in 10-cm poly-D-lysine (PDL)-coated tissue culture dishes. Cells were transfected for 24 hours with Turbofect reagent (Thermo Scientific) at a concentration of 2 μg for single plasmid transfections and 2 μg each of pCMV6 construct for co-transfections of TMEM163 and individual ZNT proteins tagged with either Myc-DDK or HA peptide (to equalize the plasmid concentrations across experiments). Twenty-four hours posttransfection, the cells were lysed with 1 mL of the Pierce Co-IP lysis buffer containing 1X protease inhibitor cocktail (PIC; Thermo Scientific) and 1 mM phenylmethylsulfonyl fluoride (PMSF) (Sigma-Aldrich). The lysates were centrifuged at 14,000 rpm for 10 minutes at 4°C to pellet and remove cellular debris. We then performed dot blots of lysates to check for expression of the peptide tagged protein and qualitatively determine protein expression before performing co-IP assays. After determining the relative concentrations of total proteins in each sample, the cell lysates were diluted with lysis buffer to a volume of 500 μL and incubated with control agarose resin for 1 hour at 4°C on a rotator to remove non-specific proteins binding to the resin. After lysate pre-clearance, equal volumes of either DDK- or HA-coupled agarose beads were added to the lysates for overnight incubation at 4°C. Protein-bound beads were washed with the lysis buffer three times and eluted with the Pierce co-IP elution buffer. To validate initial co-IP results for specificity, we repeated all experiments with a modified the wash buffer by adding 500 mM of NaCl plus 1% Tween-20 to increase the stringency of the co-IP (Cuajungco *et al.*, 2006) to possibly reveal the relative strength of the interaction between the monomers. all protein eluates were reduced with 1X Bolt sample reducing agent (Thermo Scientific) and denatured with 1X NuPAGE LDS sample buffer (Thermo Scientific). For SDS-PAGE, the samples were loaded (15 μl) in two identically arranged Bolt 4-12% gradient Bis-Tris PAGE gels (Thermo Scientific) – one gel was used for downstream visualization of DDK-tagged proteins while the other gel is for HA-tagged proteins. Resolved protein samples were then transferred to nitrocellulose membranes. One blot was probed with primary anti-DDK mAb (1:2000) while the other blot with primary anti-HA pAb (1:5000). After a series of washes with Tris-buffered solution plus 0.1% Tween-20 (TBST), all blots were incubated for 1 hour at room temperature with secondary anti-mouse or anti-rabbit IR-Dye™ 800CW (1:15000). All immunoblots were subjected to a series of washes with TBST before scanning them using the Odyssey SA™ IR imaging system (LI-COR Biosciences).

To analyze endogenous protein-protein interaction, we used postmortem mouse tissues (pancreas and testis) where mouse Tmem163 protein expression overlapped with Znt1, Znt2, Znt3, and Znt4 protein (Human Protein Atlas; www.proteinatlas.org). Before setting up the co-IP trials, we conjugated the anti-TMEM163 pAb with the agarose resin using the Pierce co-IP kit (Thermo Scientific) according to the manufacturer’s recommendations. We used primary antibody conjugation as a way to minimize co-elution of immunoglobulins whose molecular weight is relatively similar to the target protein. The frozen tissues were pulverized using mortar and pestle situated on dry ice or liquid nitrogen and homogenized with T-Per lysis buffer containing 1 mM PMSF and 2X PIC. The homogenates were centrifuged at 13,000 rpm for 15 minutes at 4°C. Except for the pancreas, we added 0.5% DDM detergent in the T-Per lysis buffer to homogenize the testis tissue to help improve membrane protein solubilization. The respective tissue homogenates were pre-cleared with control agarose resin for 30 minutes at 4°C on a rotator and then incubated overnight at 4°C with anti-TMEM163 pAb coupled to agarose beads. The following day, all samples were washed three times using the Pierce co-IP lysis buffer and the samples were recovered using the Pierce co-IP elution buffer. In cases where we observed low target protein recovery, we used the Pierce protein concentrator (10 kDa MWCO) according to the manufacturer’s protocol before performing SDS-PAGE. The samples were loaded (20 μl) into a Bolt 4-12% Bis-Tris PAGE gels and the resolved proteins were blotted. Each blot was probed with the respective primary antibody as follows: anti-TMEM163 pAb (1:500), anti-Znt1 pAb (1:500), anti-Znt2 pAb (1:400), Anti-Znt3 mAb (1:500), and anti-Znt4 pAb (1:500). The antibodies against Znt1, Znt2, and Znt4 were conjugated with Alexa-Fluor-750 using the Zenon Rabbit IgG Labelling Kit (Thermo Scientific) according to the manufacturer’s recommendation. The blots were incubated with anti-rabbit IR-Dye™ 800CW (1:15000) secondary antibody and washed with TBST several times before analyzing them using the Odyssey SA™ IR scanner.

All WB images were saved as TIF files and processed using Adobe Photoshop 2022 to crop unwanted edges or empty lanes, equalize contrast levels, and straighten vertical position. We used Adobe Illustrator 2022 to compile and label images.

### 2.5 Immunocytochemistry (ICC) and Confocal microscopy analyses

To avoid potential confounding effects of a bulky fluorescence protein tag inducing mis-localization, we expressed TMEM163 and ZNT proteins with a peptide tag and performed ICC technique. HEK-293T cells were seeded at 50,000 cells per well in 12-well tissue culture plates containing PDL-coated coverslips. Cells were then transfected with pCMV6 expression constructs of TMEM163 and ZNT proteins tagged with HA or DDK. We used 1 μg of plasmid for single transfections and also 1 μg of each plasmid for co-transfections. Twenty-four hours post-transfection, the samples were washed once with 1X phosphate-buffered saline (PBS) and fixed with 4% paraformaldehyde (PFA) for 15 minutes. The cells were washed three times with 1X PBS for 5 minutes and washed once with PBT1 (1X PBS, 0.1% BSA, 5% heat-inactivated goat serum, and 0.1% Triton X-100) and then permeabilized with PBT1 for 15 minutes. The samples were incubated with either anti-DDK mouse mAb (1:800) (Millipore Sigma) or anti-HA rabbit pAb (8 μg/mL) (Millipore Sigma) overnight at 4°C. The samples were washed four times with 1X PBS and twice with PBT2 (1X PBS, 0.1% BSA, and 0.1% TritonX-100), then incubated with secondary antibodies diluted in PBT2 for 1 hour at room temperature. For anti-DDK mouse mAb, we used anti-mouse Alexa Fluor-488 (1:500; ex = 490 nm, em = 525 nm). For anti-HA rabbit pAb, we used anti-rabbit Alexa Fluor-568 (1:500; ex = 579 nm, em = 603 nm). The coverslips were then washed six times with 1X PBS. Each coverslip was mounted on a glass slide containing a drop of Prolong Diamond anti-fade reagent plus DAPI (Thermo Scientific), and the edges were sealed with a clear nail polish. Fluorescent images were captured using an Olympus FV3000 laser scanning confocal microscope. The monochromatic images were processed using Adobe Photoshop 2022 to change grayscale images to green for Alexa-488 fluorescence, red for Alexa-568 fluorescence and blue for DAPI fluorescence. All images were cropped and auto-contrasted before the three channels were merged as a single image using Photoshop. We used Adobe Illustrator to compile and label images.

### 2.6 Cell surface biotinylation

HEK-293 cells were plated at 1.5 × 10^6^ cells in PDL-coated 10-cm petri dishes. The cells were transfected with 3 μg of DNA for single plasmids and 3 μg each plasmid for dual expression of pCMV6 constructs containing TMEM163-DDK, ZNT1-HA, ZNT2-HA, ZNT3-HA, and ZNT4-HA. In one trial, we reduced the DNA concentration to 2 μg to reduce transfection-induced cytotoxicity. Twenty-four hours post-transfection, the cells were washed two times with 10 ml of ice-cold PBS-CM (1X PBS, 0.5 mM CaCl_2_, 1 mM MgCl_2_, pH 8.0) and incubated with 1 ml of freshly prepared Sulfo-NHS-LC-LC-Biotin solution (0.5 mg/ml in PBS-CM) for 30 min at 4°C. The cells were washed twice with cold 10 ml of PBS-CM containing 0.1% BSA to terminate and quench any unbound Sulfo-NHS-LC-LC-Biotin. The cells were washed once with PBS (pH 7.4) and then lysed for 1 hour at 4°C with 1 ml of lysis buffer (150 mM NaCl, 5 mM EDTA, 1% Nonidet P-40, 50 mM Tris, pH 7.5) supplemented with 1X PIC and 1 mM PMSF. The cell lysates were centrifuged at 14,000 rpm for 10 min at 4°C. We took an aliquot (1-5%) from each sample to measure total protein concentration using the BCA assay, in order to normalize the total protein concentration of each cell lysate. We equilibrated the Neutravidin beads in lysis buffer, added 150 μl of the beads to all normalized protein samples, and incubated the samples rotating overnight at 4°C. The samples were washed five times with lysis buffer, incubated in 65 μl elution buffer (2X Bolt LDS buffer) for 30 minutes at 37°C, and eluted by low speed centrifugation. The samples were immunoblotted with primary anti-DDK mAb or anti-HA pAb and visualized with secondary anti-mouse or anti-rabbit IR-Dye 800CW, respectively. The blots were then scanned using the Odyssey SA™ IR imager. The images were processed using Adobe Photoshop 2022 and the figure was created using Adobe Illustrator 2022.

### 2.7 Spectrofluorometric zinc flux assay

To determine zinc flux in cells transfected with pBI constructs, we seeded HeLa cells at 10,000 cells per well on a 96-well culture plate treated with PDL. Untransfected cells served as negative control. We used cell membrane impermeant Fluozin-3 (FZ3, Kd ~15 nM; ex = 494 nm, em = 516 nm) and cell membrane permeant Newport Green (NG, Kd ~1 μM; ex = 505 nm, em = 535 nm) dyes to assay changes in extracellular and intracellular zinc levels, respectively, as described in our protocol paper (Ali and Cuajungco, 2020). In brief, we used a standard amount of 150 ng of DNA constructs per well using Turbofect lipid reagent. Twenty-four hours post transfection, we proceeded with either the FZ3 or NG assay.

For FZ3 assay, we used Kreb’s Ringers Bicarbonate buffer (14.99 mM sodium bicarbonate, 119.8 mM NaCl, 4.56 mM KCl, 0.49 mM MgCl_2_, 0.70 mM sodium phosphate dibasic, 1.30 mM sodium phosphate monobasic, 10 mM glucose, pH 7.4) supplemented with 2 mM Glutamax and 1 mM sodium pyruvate. The treatment group was exposed to zinc chloride (ZnCl_2_; 200 μM) plus zinc pyrithione (20 μM) and the cells were placed in a standard humidified 37°C incubator with 5% CO_2_ for 10 minutes.

For NG assay, we used standard HEPES buffer (10 mM HEPES, 135 mM NaCl, 5 mM KCl, 1mM CaCl_2_, 5 mM glucose, pH 7.4) supplemented with 2 mM Glutamax and 1 mM sodium pyruvate. The cells were first incubated with NG (5 μM) for 20 minutes inside the 37°C cell culture incubator, washed twice with HEPES buffer, and placed at room temperature for 30 minutes to allow for NG ester cleavage. The treatment group was exposed to zinc chloride (ZnCl_2_; 100 μM) plus zinc pyrithione (10 μM) and the cells were placed in the 37°C cell incubator for 20 minutes.

After pre-loading the treatment group with zinc, the cells for both assays were washed five times with the respective assay buffer using an automated BioTek 405 TS microplate washer (BioTek Instruments, Winooski, VT). For the final aspiration after the last wash, each well was injected with 100 μL of the respective assay buffer. The plate was transferred into the BioTek Cytation 5 plate reader. For the FZ3 assay, each well was injected with the membrane impermeable FZ3 dye to a final concentration of 1 μM and kinetic readings were done for each well every minute for 25 minutes. For the NG assay, the plates were placed in the plate reader after the last wash and kinetic readings for each well were done every minute for 45 minutes. In general, the averaged background relative fluorescence unit (RFU) values from non-treatment group (cells transfected but not pre-loaded with zinc) were subtracted from RFU values of treatment group pre-loaded with zinc.

### 2.8 Statistical Analysis

Spectrofluorometric data were evaluated for statistical significance using GraphPad Prism version 9 software (La Jolla, CA). RFU values from Fluozin-3 trials were analyzed using one-way analysis of variance (ANOVA) with repeated measures followed by Tukey’s *post-hoc* multiple comparisons test (control versus treatment). RFU values from Newport Green trials were analyzed using The significance level was set at *p* < 0.05.

## 3. RESULTS

We initially performed co-IP experiments using the sample wash buffer from the Pierce co-IP kit, which is a standard buffer. However, to ensure that the co-IP results were specific, we subsequently used a highly stringent wash buffer in subsequent experiments to show the extent of protein-protein interactions between TMEM163 dimer and TMEM163/ZNT heterodimers. As expected, co-IP experiments showed that human TMEM163 formed a dimer with itself (**Fig. 1**), which served as a control for all other experiments where we co-expressed TMEM163 with ZNT proteins. We then found that ZNT1-HA or ZNT2-HA co-expressed with TMEM163-DDK resulted in co-elution of TMEM163-DDK when ZNT1-HA or ZNT2-HA was immunoprecipitated (**Figs. 1A-1B**). Co-expression of ZNT3-HA with TMEM163-DDK also showed interaction between the two proteins as shown by the co-eluted TMEM163-DDK band upon immunoprecipitation of ZNT3-HA (**Fig. 1C-1D**). Likewise, co-expression of ZNT4-DDK with TMEM163-HA resulted in co-elution of ZNT4-DDK after TMEM163-HA was immunoprecipitated (**Fig. 1C-1D**). Interestingly, the highly stringent washing of the co-IP samples revealed that the heterodimeric binding between TMEM163 and ZNT proteins was not as physically strong as the homodimer interaction between TMEM163 monomers (e.g., compare the WB band elution profiles of all TMEM163 co-IP with itself versus those WB elution bands of TMEM163 co-IP with specific ZNT proteins).

**Figure 1.**
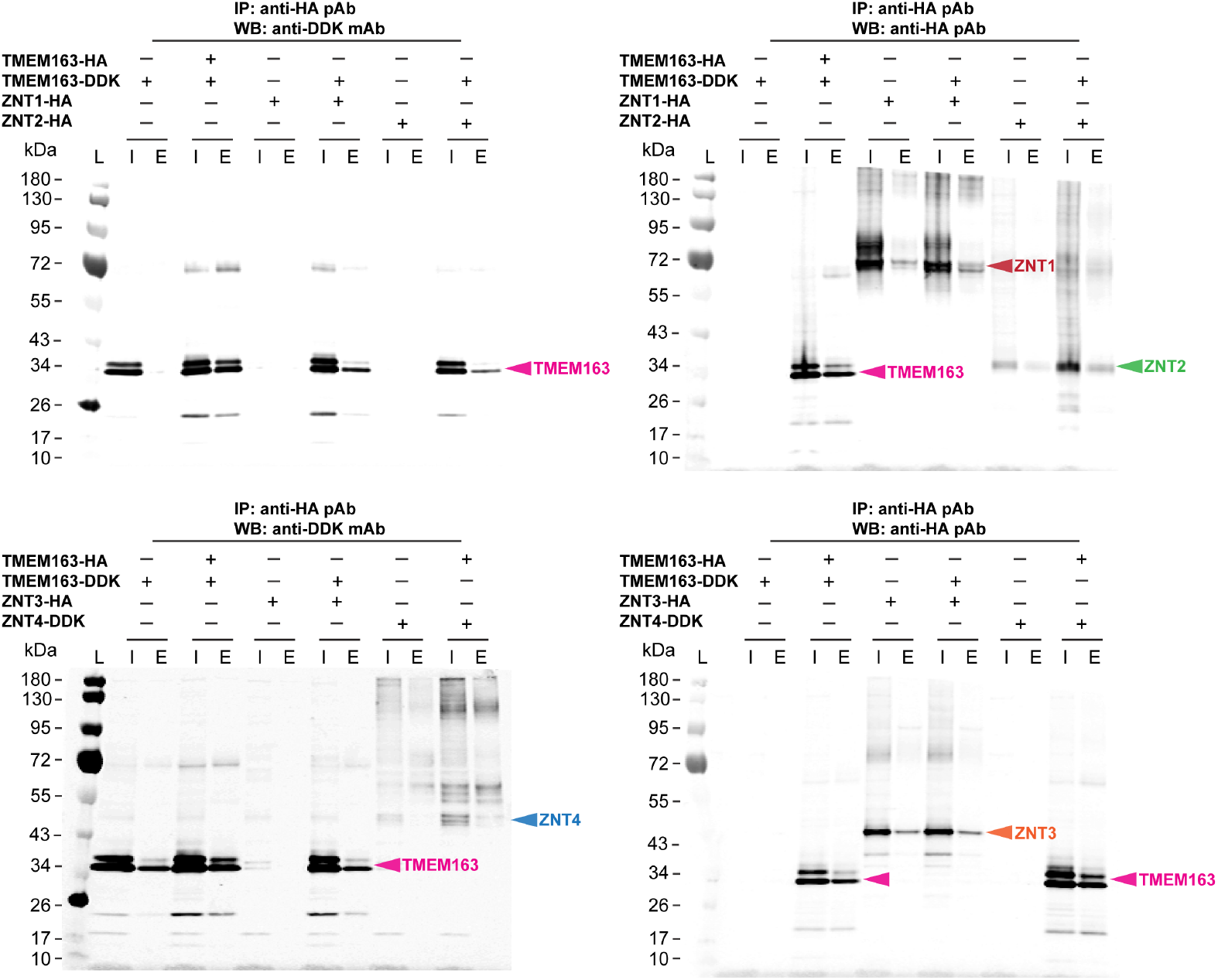
Co-immunoprecipitation assays of heterologously expressed TMEM163, ZNT1, ZNT2, ZNT3, and ZNT4 proteins. Co-expressions of TMEM163-DDK with ZNT1-HA, ZNT2-HA, and TMEM163-HA show heterodimerization as indicated by their respective eluted bands upon immunoblotting with **A**) anti-DDK monoclonal antibody (mAb) and **B**) anti-HA polyclonal antibody (pAb). Co-expressions of TMEM163-DDK with ZNT2-HA, and TMEM163-HA with ZNT4-DDK both show heterodimerization as indicated by their respective eluted bands upon immunoblotting with **C**) anti-DDK mAb and **D**) anti-HA pAb. Expression of TMEM163-HA or TMEM163-DDK in all trials serves as a positive (dimerization) control with itself, while single expressions of each respective protein serve as negative controls. L, protein ladder; I, input/cell lysate; E, elution; +, expressed; –, not expressed. Predicted molecular weight (MW) of human proteins: TMEM163 = 31.5 kDa, ZNT1 = 55.3-63.0 kDa, ZNT2 variant 2 = 35 kDa, ZNT3 = 41.8 kDa, and ZNT4 = 47.4 kDa. The images are representative of N ≥ 6 independent experiments.

In a separate set of experiments, we performed co-IP of TMEM163 with each of the ZNT protein examined that included control agarose beads as an additional approach to guarantee the specificity of protein binding to the antibody-conjugated beads. We confirmed that the interactions are specific and did not observe non-specific binding in the absence of the primary antibody (**Fig. S2**). To further corroborate our findings, we performed native co-IP assays using mouse tissues. We had *a priori* knowledge that Tmem163 is expressed in the pancreas, and we then checked which tissues express Znt1, Znt2, Znt3 and Znt4 proteins using the Human Protein Atlas database. We found that Znt2 and Znt4 are present in the pancreas, while Znt1 and Znt3 are present in testis. We tested antibodies against mouse Tmem163, Znt1, Znt2, Znt3 and Znt4 proteins whether they will cross-react to their human protein counterparts by using cell lysates over-expressing each of the human protein. Upon determining antibody cross-reactivity, we used the respective human protein over-expression (OE) cell lysates as a positive control to identify endogenous mouse Tmem163 and Znt proteins. We used an anti-TMEM163 polyclonal antibody (pAb) that we previously validated (Cuajungco *et al.*, 2014; Sanchez *et al.*, 2019) to co-IP endogenous Tmem163, Znt2 and Znt3 proteins from pancreatic tissue homogenates. Similarly, we used the same anti-TMEM163 pAb to co-IP Znt1 and Znt3 proteins from testis tissue homogenates. WB analysis showed that native Tmem163 proteins co-eluted with itself upon co-IP (**Fig. 2**). Endogenous Znt1, Znt2, Znt3, and Znt4 proteins also co-eluted with Tmem163 as evidenced by WB probed with antibodies against the respective Znt proteins (**Fig. 2**). Taken together, heterologous expression and native co-IP experiments indicate that TMEM163 heterodimerizes with ZNT1-ZNT4 proteins.

**Figure 2.**
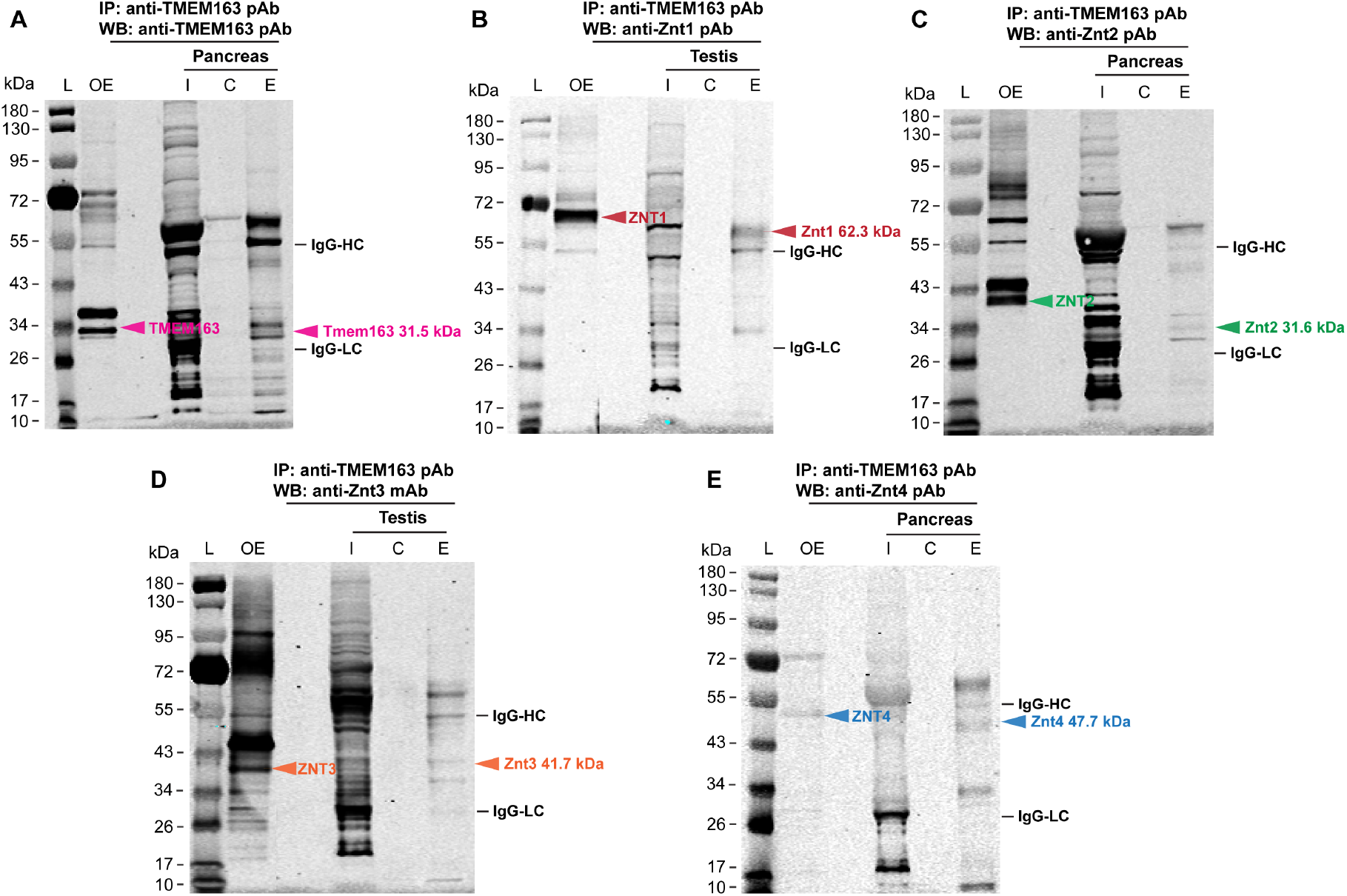
Endogenous mouse Tmem163 protein interacts with Znt1, Znt2, Znt3, and Znt4 proteins. **A**) Native co-IP Western blot (WB) of mouse Tmem163 protein from pancreas shows co-elution with itself using anti-TMEM163 polyclonal antibody (pAb). **B**) Native co-IP WB of mouse Znt1 protein from testis co-elutes with Tmem163 as shown by anti-Znt1 pAb. **C**) Native co-IP WB of mouse Znt2 protein from pancreas co-elutes with Tmem163 as shown by anti-Znt2 pAb. **D**) Native co-IP WB of mouse Znt3 protein from testis co-elutes with Tmem163 as shown by anti-Znt3 monoclonal antibody (mAb). **E**) Native co-IP WB of mouse Znt4 protein from pancreas coelutes with Tmem163 as shown by anti-Znt4 pAb. Cell lysates of over-expressed (OE) human TMEM163, ZNT1, ZNT2, ZNT3, and ZNT4 proteins serve as positive expression control while the agarose control beads serve as negative control for their respective immunoblots. Predicted molecular weight (MW) of mouse proteins: Tmem163 = 31.5 kDa, Znt1 = 54.6-62.3 kDa, Znt2 = 31.6 kDa, Znt3 = 41.7 kDa, and Znt4 = 47.7 kDa. Predicted MW of OE human proteins: TMEM163 = 31.5 kDa, ZNT1 = 55.3-63.0 kDa, ZNT2 variant 1 = 40.4 kDa, ZNT3 = 41.8 kDa, and ZNT4 = 47.4 kDa. IgG-HC, immunoglobulin-G heavy chain (MW @ 55 kDa); IgG-LC, immunoglobulin-G light chain (MW @ 23 kDa). L, protein ladder; I, input/cell lysate; C, control resin; E, elution. The images are representative of N ≥ 3 independent experiments.

We then investigated the functional significance of TMEM163 interacting with each of the four ZNT proteins using Fluozin-3 and Newport Green zinc flux assays (Ali and Cuajungco, 2020). To accomplish this study, we heterologously expressed pBI constructs with single ORFs (TMEM163, ZNT1, ZNT2, ZNT3, and ZNT4) and pBI constructs with dual ORFs (TMEM163+ZNT1, TMEM163+ZNT2, TMEM163+ZNT3, and TMEM163+ZNT4) in their corresponding combinations. Our results confirmed our previous report that TMEM163 is a zinc effluxer whereby its cellular expression significantly reduced intracellular zinc levels in comparison to untransfected control cells (p < 0.0001, ANOVA, Tukey’s multiple comparisons test) (**Fig. 3**). Further, the zinc efflux profiles between TMEM163 homodimers and TMEM163/ZNT heterodimers were consistently different in which the TMEM163/ZNT heterodimers appear to extrude zinc slightly better than TMEM163 homodimer. Noteworthy, we found that co-expression of TMEM163 with ZNT1 resulted in an even more significant enhancement of zinc efflux activity over time for TMEM163/ZNT1 heterodimer compared with TMEM163 homodimer as demonstrated by both zinc flux assays (**Fig. 3**).

**Figure 3.**
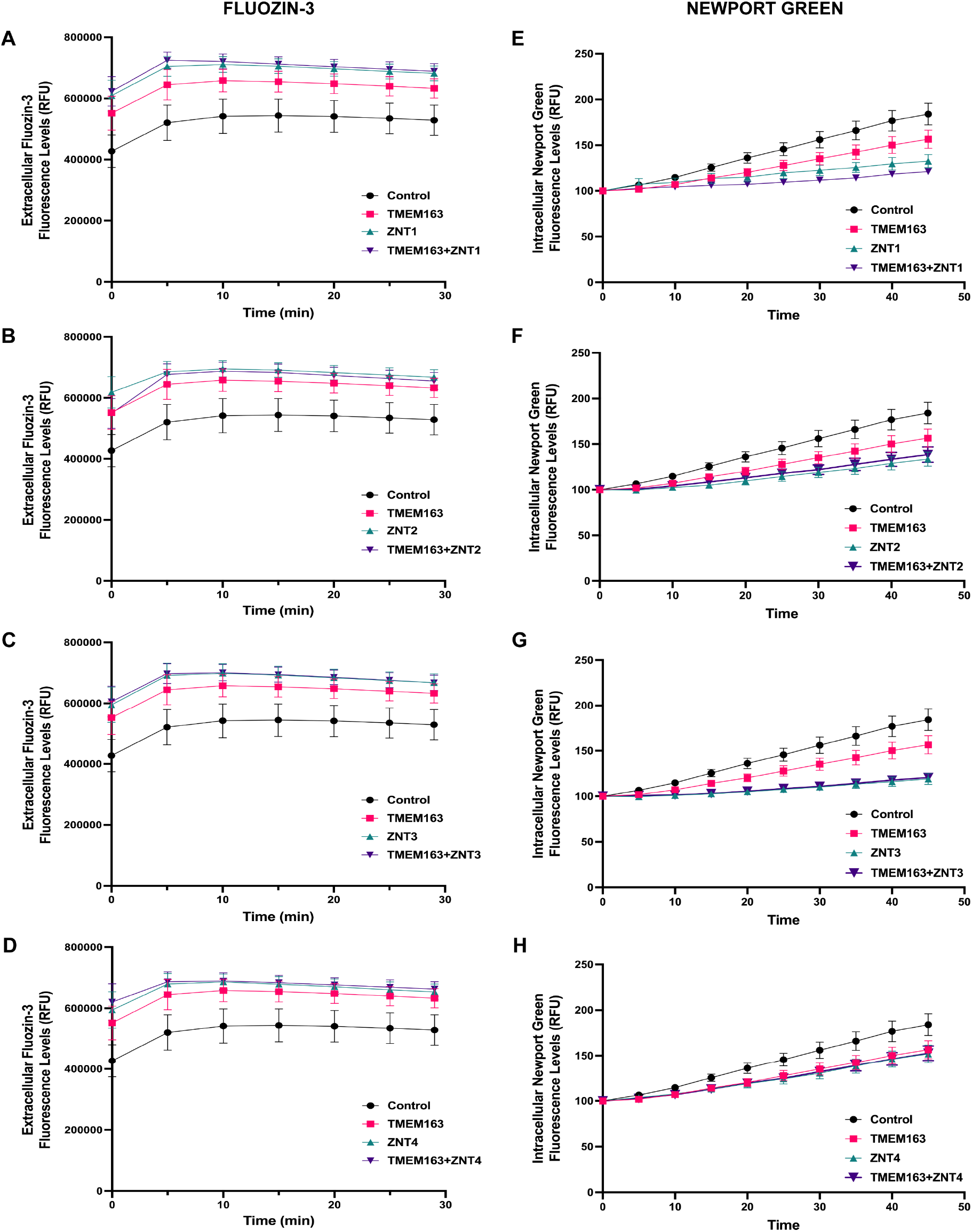
TMEM163 and ZNT proteins efflux zinc. Zinc flux assays using the cell membrane impermeant Fluozin-3 dye (**A-D**) and cell membrane permeant Newport Green dye (**E-G**) upon single expression and co-expression of TMEM163, ZNT1, ZNT2, ZNT3, and ZNT4. Heterologous expression of TMEM163 and ZNTx (where x denotes 1, 2, 3 or 4) show significant zinc extrusion when compared to untransfected control (p < 0.0001, Tukey’s multiple comparisons test). Data shown are 5-minute intervals and represented as mean ± SEM (N ≥ 4 independent trials). RFU, relative fluorescence unit.

Golan and colleagues previously reported that ZNT1, ZNT2, ZNT3 and ZNT4 monomers form heterodimers (Golan *et al.*, 2015). The group found that ZNT1-ZNT3 heterodimers were mostly localized within intracellular vesicles, which effectively reduced the PM localization of ZNT1 protein, while heterodimerization of ZNT2 or ZNT4 with ZNT1 increased the PM localization of both ZNT2 and ZNT4 (Golan *et al.*, 2015). The group’s observations suggest that certain ZNT monomers influence their membrane localization. To investigate whether the interaction of TMEM163 with ZNT proteins also impacts their subcellular localization and vice versa, we performed ICC experiments upon co-expression of TMEM163 with ZNT proteins. The ICC approach is also a way to further validate our co-IP data since we expected these proteins should co-localize together if they form heterodimers in their particular cellular sites. We also heterologously expressed TMEM163, ZNT1, ZNT2, ZNT3, and ZNT4 proteins individually in cells to visually compare their subcellular localization with dual TMEM163/ZNT protein expression. Confocal microscopy showed that TMEM163 protein is localized within the cell periphery and intracellular compartments (**Fig. S3**), which is consistent with our previous reports (Cuajungco *et al.*, 2014; Sanchez *et al.*, 2019). ZNT1 exhibited a similar localization as TMEM163, while ZNT2, ZNT3, and ZNT4 showed punctate distribution patterns in cells (**Fig. S3**). These results are consistent with previous observations for all ZNT proteins studied (Lopez and Kelleher, 2009; Salazar *et al.*, 2009; McCormick and Kelleher, 2012; Golan *et al.*, 2015; Nishito and Kambe, 2019). Meanwhile, co-expression of TMEM163 with each of the ZNT protein revealed partial co-localization of the corresponding TMEM163/ZNT heterodimers (**Fig. 4**). The localization patterns of TMEM163 and ZNT proteins did not show any prominent alteration in their subcellular localization when expressed alone (e.g., compare **Fig. S3** with **Fig. 4**). To examine the cell surface expression of TMEM163 and ZNT proteins, we biotinylated heterologously expressed proteins containing either DDK or HA peptide tag. We found that the PM localization of TMEM163 did not change significantly when it was co-expressed with ZNT1 or ZNT4, but a more obvious difference in cell surface expression of TMEM163 was evident in the presence of ZNT2 or ZNT3, which demonstrated relatively lighter protein band intensities (**Fig. 5A**). On the other hand, the co-expression of TMEM163 with each of ZNT protein seemed to modestly change their cell surface expression relative to their single expression profiles (**Fig. 5B**). It is worth noting that the ZNT1 protein that is detected on the cell surface appeared to have a darker upper band compared to its lower band upon co-expression with TMEM163, suggesting that there were relatively more ZNT1 isoforms with post-translational modification (PTM) detected in the PM. ZNT1 has been reported to be N-glycosylated, which makes it appear as double band on WB (Nishito and Kambe, 2019). The upper band or N-glycosylated isoform of ZNT1 has an apparent molecular weight (MW) of 75 kDa while its non-glycosylated isoform (lower band) is around 63 kDa even though its theoretical MW is 55.3 kDa (Nishito and Kambe, 2019). It is interesting to note that when ZNT1 was co-eluted with TMEM163, its WB band profile displayed a bias towards the non-PTM (non-glycosylated) isoform (see **Fig. 1** and **Fig. S2**). This observation is similar to the TMEM163 WB band profile that is detected on the cell surface, which appeared to be non-PTM isoforms since the lower band was consistently darker than the higher band (see **Fig. 1** and **Fig. 5A**). We attributed the higher WB band of TMEM163 protein as PTM based on our *in silico* analysis using the Phosphosite online software (www.phosphosite.org) predicting six phosphorylation sites (five Serine residues and one Threonine residue) within its N-terminus (NT) region (data not shown). Finally, the detection of ZNT2, ZNT3, and ZNT4 on the cell surface upon biotinylation may be a likely artifact of their over-expression in cells since ZNT1 is only expected to be localized within the PM (Nishito and Kambe, 2019). Notwithstanding, our data supported the observation that certain protomers of the ZNT protein family form functional heterodimers with each other (Salazar *et al.*, 2009; Golan *et al.*, 2015; Zhao *et al.*, 2016) but that this unique group also now includes TMEM163 protein. Indeed, the predicted secondary structure of TMEM163 and ZNT proteins (**Fig. S4**) adds credence to the results of this study, since we believe that monomers with similar structural features such as six transmembrane domains will likely interact as a dimer.

**Figure 4.**
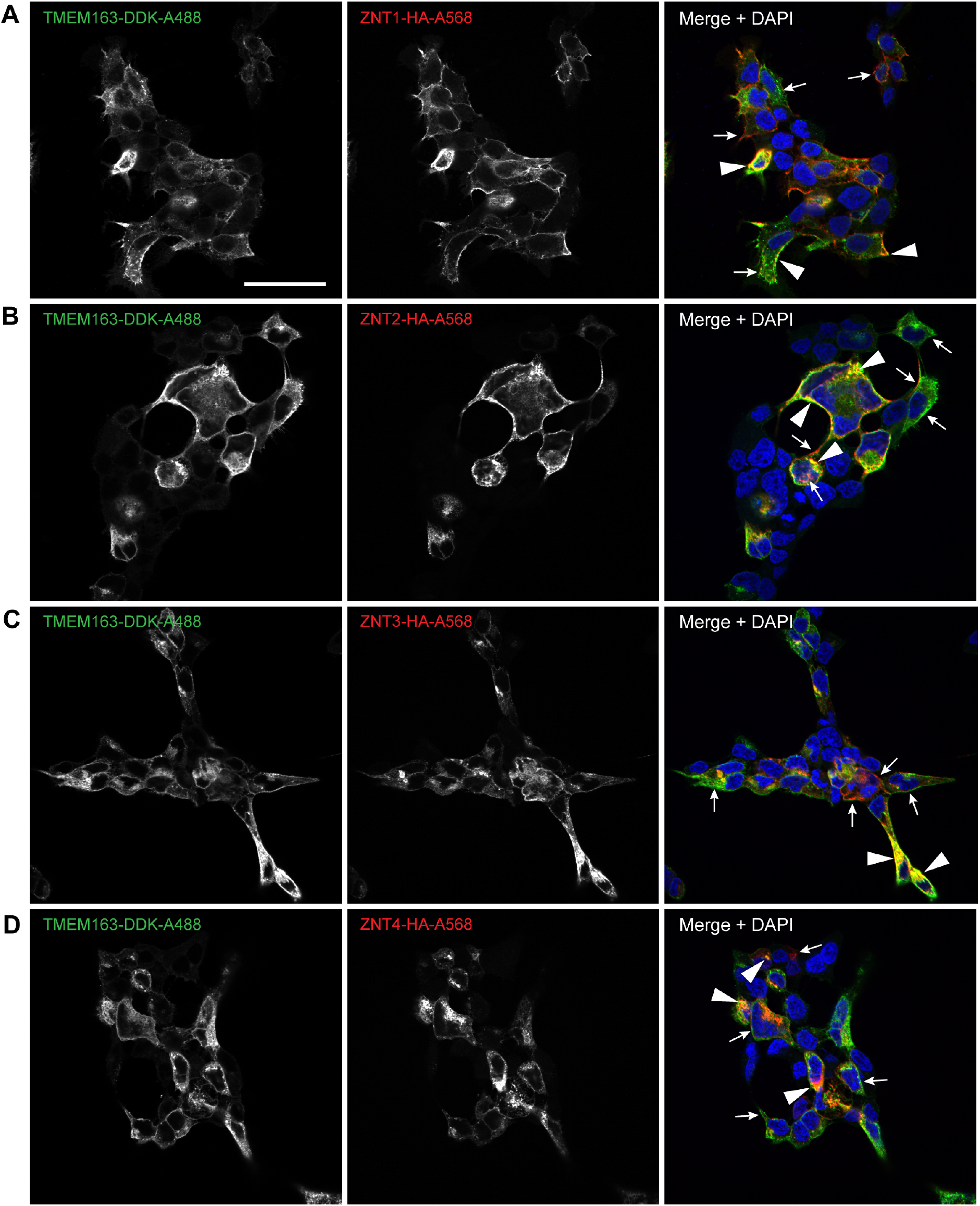
Immunocytochemistry of TMEM163 heterologously co-expressed with ZNT proteins. Representative confocal images of cells co-transfected with TMEM163 and **A**) ZNT1-HA, **B**) ZNT2-HA, **C**) ZNT3-HA, and **D**) ZNT4-HA with TMEM163-DDK. The DDK peptide was detected by anti-DDK mAb and visualized with anti-mouse secondary antibody conjugated with Alexa-488 (green). The HA peptide was detected by anti-HA pAb and visualized with anti-rabbit secondary antibody conjugated with Alexa-568 (red). DAPI stains the nuclei blue. Arrow indicates co-localization of TMEM163 and ZNT protein heterodimers within their respective cellular sites. Arrowhead indicates non-overlapping subcellular localization of protein homodimers. Scale bar: 100 μm.

**Figure 5.**
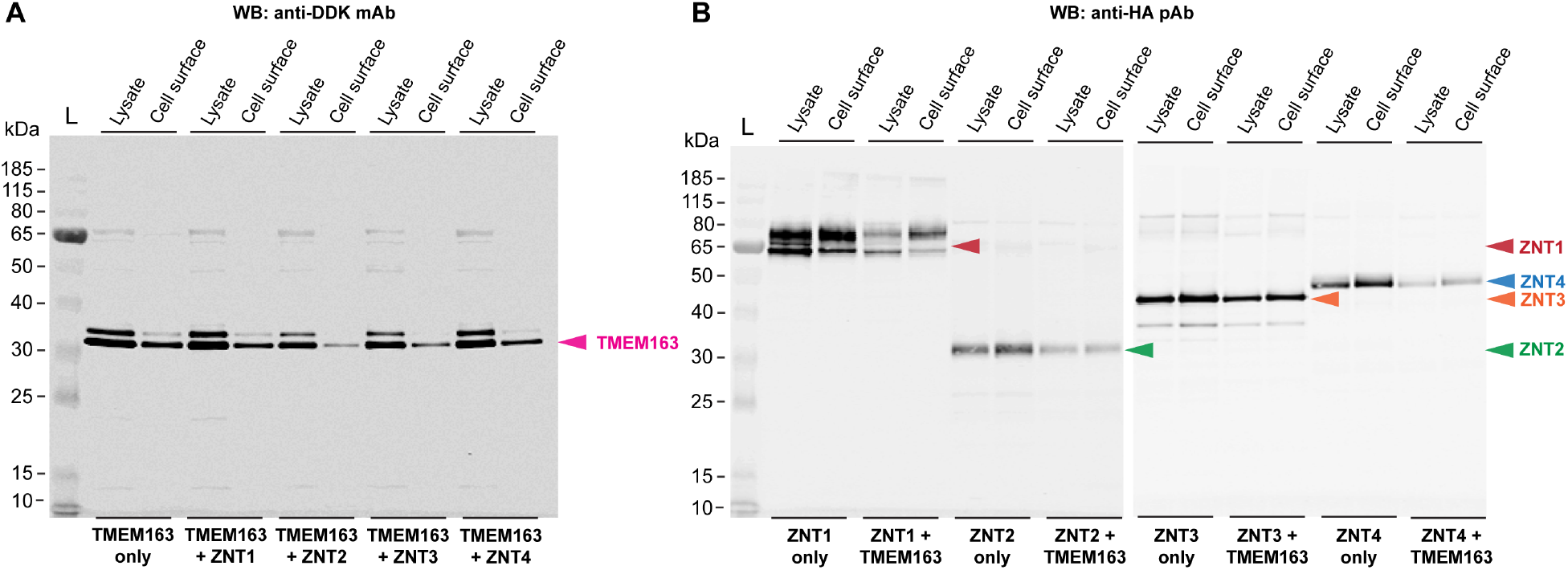
Cell surface expression of TMEM163 and ZNT proteins. **A**) TMEM163 co-expression with ZNT1, ZNT2, ZNT3, and ZNT4 proteins. A slight decrease in PM localization of TMEM163 is observed in the presence of ZNT2 and ZNT3, but not ZNT1 or ZNT4. **B**) Cell surface expression of ZNT1, ZNT2, ZNT3, and ZNT4 proteins show a modest decrease in the PM localization upon co-expression with TMEM163 relative to their single expression. Predicted MW of human proteins: TMEM163 = 31.5 kDa, ZNT1 = 55.3-63.0 kDa, ZNT2 variant 2 = 35 kDa, ZNT3 = 41.8 kDa, and ZNT4 = 47.4 kDa.

## 4. DISCUSSION

TMEM163 is a zinc transporter that shares evolutionary conservation with certain members of the CDF protein superfamily (Sanchez *et al.*, 2019; Styrpejko and Cuajungco, 2021). Our current study further confirms that TMEM163 extrudes intracellular zinc as a dimer or heterodimer with distinct ZNT proteins. Thus, TMEM163 is a likely *bona fide* member of the SLC30/ZNT zinc efflux transporters as evidenced by its functional interaction with other ZNT proteins. The highly stringent washing in our co-IP experiments showed that the physical binding between TMEM163 and ZNT monomers is relatively weak compared to TMEM163 homodimer. Such difference suggests that the interaction domain between TMEM163 and ZNT proteins may not be the same as the dimerization domain. This conjecture is potentially bolstered by the previous observation that the dimerization of ZNT3 protomer is mediated by covalently linked dityrosine amino acid residues (Salazar *et al.*, 2009), but that the same tyrosine residues responsible for ZNT3 dimerization are not critical for ZNT3/ZNT4 or ZNT3/ZNT10 heterodimerization (Zhao *et al.*, 2016). Future work should address the possibility that the dimerization and heterodimerization domains of TMEM163 may parallel that of the ZNT3 protein. Meanwhile, the presence of some non-specific bands in our native co-IP blots is due to the polyclonal nature of the antibodies used in the study, but the inclusion of positive over-expression control and the use of IgG heavy/light chains as landmarks, coupled with the molecular weight marker, helped us identify each of the endogenous mouse Znt protein co-eluting with Tmem163 protein. In addition, confocal imaging also provided evidence that heterodimers of TMEM163 and ZNT proteins exist in their respective cellular sites, albeit they only partially co-localized. This observation suggests that TMEM163 or ZNT homodimers may function differently that their heterodimer equivalents. Indeed, this is exactly what we observed using two different zinc flux assays (**Fig. 4**). The magnitude of zinc efflux activity in cells expressing TMEM163 alone varies when compared with cells expressing both TMEM163 and each of the ZNT protein. What is intriguing, however, is the noticeable enhancement of zinc efflux activity of TMEM163/ZNT1 heterodimer compared with TMEM163 homodimer. The relative increase in the N-glycosylated isoform of ZNT1 (Nishito and Kambe, 2019) on the cell surface when co-expressed with TMEM163 (**Fig. 5B**) could explain such difference in efflux activity since both proteins have been reported to be present in the PM (Cuajungco *et al.*, 2014; Nishito and Kambe, 2019) while ZNT2, ZNT3, and ZNT4 are mostly detected within intracellular compartments (Lopez and Kelleher, 2009; Salazar *et al.*, 2009; McCormick and Kelleher, 2012; Golan *et al.*, 2015). Overall, diversifying the protein composition of zinc effluxers in cells may be a crucial way for them to efficiently eliminate excess cytoplasmic zinc when such need arises as in the case of cells expressing TMEM163/ZNT1 heterodimers exposed to high levels of exogenous zinc. This phenomenon may also explain why many cell types have redundant expressions of influx and efflux zinc transporters. Although the zinc efflux activity of TMEM163/ZNT3 heterodimer showed a similar synergy for effluxing zinc relative to TMEM163 homodimer for the Newport Green assay, the results from the Fluozin-3 assay did not mirror this effect possibly because of higher errors obtained in the experiments. It may be that the Fluozin-3 data for TMEM163/ZNT3 heterodimer accurately reflect its efflux function since our confocal imaging and cell surface biotinylation observations indicate that relatively less TMEM163 protein is present in the PM when it is co-expressed with ZNT3 (**Figs. 4 and 5**). On the other hand, it may be possible that the TMEM163/ZNT3 heterodimer is indeed more efficient in effluxing or shuttling zinc into the lumen of various membrane compartments where the heterodimers are present. Additional research is needed to explain this discrepancy, especially that the ensuing reduction of Newport Green dye fluorescence may indicate a change in cytoplasmic zinc levels due to zinc transport in cellular compartments and extracellular space while the increase in Fluozin-3 fluorescence is mainly indicative of extracellular zinc elevation caused by cytoplasmic zinc extrusion into the extracellular milieu.

As mentioned earlier, the WB band profile typically observed for TMEM163 protein appears as double bands. We surmised that the top band is a product of PTM not only because TMEM163 has six predicted phosphorylation sites in its NT region but that our previous findings showed that deletion of at least 42 amino acid residues in its NT region eliminates one of the double bands (Cuajungco *et al.*, 2014). Furthermore, a TMEM163 protein variant with serine substituted with arginine at position 61 (S61R) resolves slightly faster and appears lower than the wildtype double bands on WB (Sanchez *et al.*, 2019). Relevant to a possible PTM of TMEM163 is the observation that its WB elution profiles upon co-IP (**Fig. 1**) and cell surface biotinylation with or without co-expressed ZNT protomers (**Fig. 5**), appear to be mostly non-PTM protein isoforms (darker lower band relative to its upper band). On the other hand, the WB elution profile of TMEM163 as a homodimer shows similar intensities between the upper and lower bands, suggesting that its dimerization domain may be subject to PTM (e.g., phosphorylation). These findings also imply that the interaction between TMEM163 and ZNT monomers may be occurring before TMEM163 undergoes PTM or that PTM favors the formation of TMEM163 homodimers. The former reasoning is supported by a similar observation that non-glycosylated ZNT1 isoform (lower band of approximately 63 kDa on WB) mainly co-eluted with TMEM163 protein (see **Fig. 1** and **Fig. S2**). It would be interesting to determine in future investigations whether or not the predicted phosphorylation status of TMEM163 at the NT region influences its homodimerization, its heterodimerization with certain ZNT protomers, and overall zinc efflux function.

In conclusion, we discovered that TMEM163 is a promiscuous protein that interacts not only with TRPML1 and P2X3/P2X4 receptor ion channels (Cuajungco *et al.*, 2014; Salm *et al.*, 2020) and BLOC-1 cargo trafficking protein (Yuan *et al.*, 2021), but also heterodimerizes with functionally related zinc efflux transporters ZNT1, ZNT2, ZNT3 and ZNT4. The physiological significance of the interaction between TMEM163 and ZNT proteins not only provides a diverse kind of zinc transporters, but also adds an extra layer of redundancy in tightly regulating zinc levels and protecting specific cells or tissues from intracellular zinc overload.

## ACKNOWLEDGMENTS

We are grateful to Nimrah Ashfaq and Steve Karl for their technical support. We also thank Dr. Shannon Kelleher (University of Massachusetts, Amherst) for providing the ZnT2 clone, and Dr. Robert Palmiter (University of Washington, Seattle) for providing the ZnT3 clone.

## FUNDING

This work was supported by the National Institutes of Health (NIH), National Institute of Neurological Diseases and Stroke AREA grant 2R15 NS0101594-01 and partly funded by 1R03 NS123728-01 to MPC. The content of this paper is solely the responsibility of the authors and does not necessarily represent the official views of the National Institutes of Health.

## DECLARATION OF COMPETING INTERESTS

The authors declare no competing financial interests or conflicts that may affect the contents of this research article.

## Notes

### Competing Interest Statement

The authors have declared no competing interest.

