## Supplemental Data for "Transmembrane 163 (TMEM163) protein interacts with specific mammalian SLC30 zinc efflux transporter family members"

### SUPPORTING INFORMATION

**TABLE S1. PCR primer sets.** List of In-Fusion (IF) cloning and restriction enzyme (RE) subcloning primers used in this study. In-Fusion primers were designed using the Takara Bio online In-Fusion cloning primer design tools for all pBI constructs and two pCMV6 vectors (ZNT2-HA and ZNT4-HA). For the IF primers designed for pCMV6, we included restriction sites (*Sgf I* and *Mlu I*) for subcloning purpose in case that IF cloning fails. Primers with restriction enzyme sites were used to clone ZNT1 and 3 into the pCMV6 vector with HA or DDK tag using *Sgf I* and *Mlu I* restriction enzymes.

| Gene/Plasmid | Primer Sequence |
|---|---|
| **Human *TMEM163*** pBI construct (MCS1) | IF Forward: 5'-TCT AGA GAC CAT GGA GCC GGC CGC GGG C-3'<br>IF Reverse: 5'-GAA TTC TCT CAA ACA TCT CGT AGT G-3' |
| **Human *ZNT1*** pBI construct (MCS1) | IF Forward: 5'-TAG TCA GCT GAC GCG ACC ATG GGG TGT TGG GGT CGG-3'<br>IF Reverse: 5'-CCG CGC TAG CAC GCG TCA CAA AGA TGA TTC AGG TTG-3' |
| pCMV6 construct | RE Forward: 5'-AAA AAA GCG ATC GCA CCA TGG GGT GTT GGG GTC GGA AC-3'<br>RE Reverse: 5'-AAA AAA ACG CGT CAA AGA TGA TTC AGG TTG TTT GTT TG-3' |
| **Human *ZNT2*** pBI construct (MCS1) | IF Forward: 5'-TAG TCA GCT GAC GCG ACC ATG GAG GCC AAG GAG AAG CAG-3' |

| | |
|---|---|
| | IF Reverse: 5’-CCG CGC TAG CAC GCG TCA GTC TGA GGG GCC CTG-3’ |
| pCMV6 construct | IF Forward: 5’-GGA GAT CTG CCG CCG CGA TCG CAC CAT GGA GGC CAA GGA GAA GCA GCA T-3’<br>IF Reverse: 5’-GAG CGG CCG CGT ACG CGT GTC TGA GGG GCC CTG GCA TGC-3’ |
| **Human *ZNT3***<br>pBI construct (MCS1) | IF Forward: 5’-TAG TCA GCT GAC GCG ACC ATG GAG CCC TCT CCA GCC G-3’<br>IF Reverse: 5’-CCG CGC TAG CAC GCG TCA GGC TTG GGG GGG TTC-3’ |
| pCMV6 construct | RE Forward: 5’-AAA AAA GCG ATC GCA CCA TGG AGC CCT CTC CAG CCG-3’<br>RE Reverse: 5’-AAA AAA ACG CGT GGC TTG GGG GGG TTC CTG-3’ |
| **Human *ZNT4***<br>pBI construct (MCS1) | IF Forward: 5’-TAG TCA GCT GAC GCG ACC ATG GCC GGC TCT GGC GCG-3’<br>IF Reverse: 5’-CCG CGC TAG CAC GCG TTA GGG ACT AGA ACT CTG AC-3’ |
| pCMV6 construct | IF Forward: 5’-GGA GAT CTG CCG CCG CGA TCG CAC CAT GGC CGG CTC TGG CGC GTG GAA G-3’<br>IF Reverse: 5’-GAG CGG CCG CGT ACG CGT GGG ACT AGA ACT CTG ACA ATT TG-3’ |

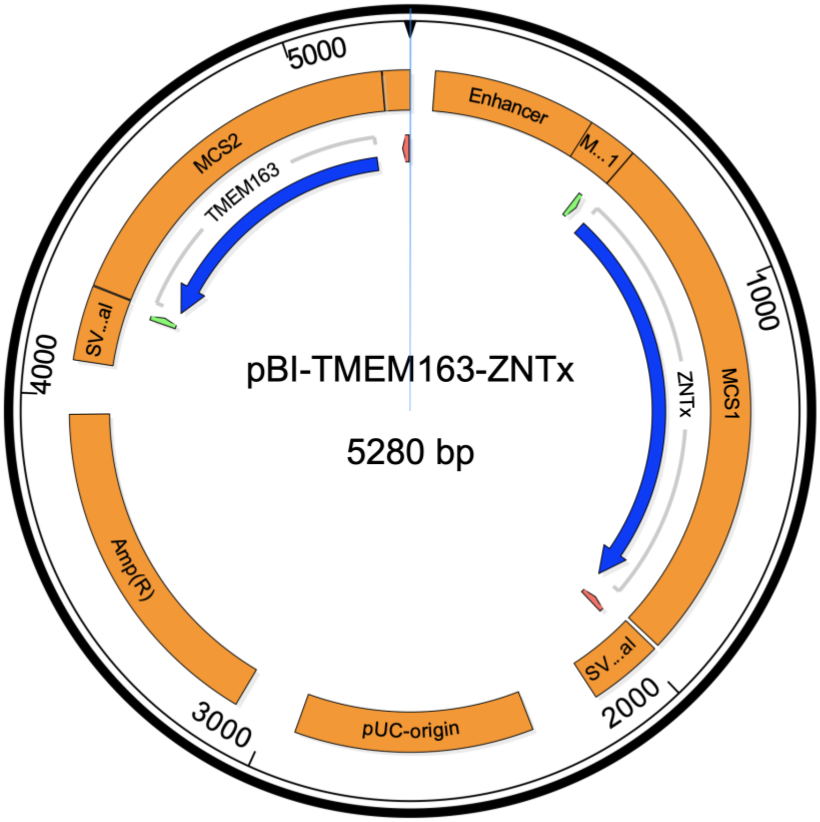


**Figure S1**. **Schematic diagram of the pBI dual expression vector.** Open reading frame (ORF) of TMEM163 was cloned into the MCS2 while ORF of each ZNTx (where x denotes 1, 2, 3, or 4) was cloned into MCS1 as described in the Materials and Methods section. Single expression constructs use the same configuration but leaving one of the other MCS empty. The pBI vector was purchased from Takara Bio, USA (Mountain View, CA).

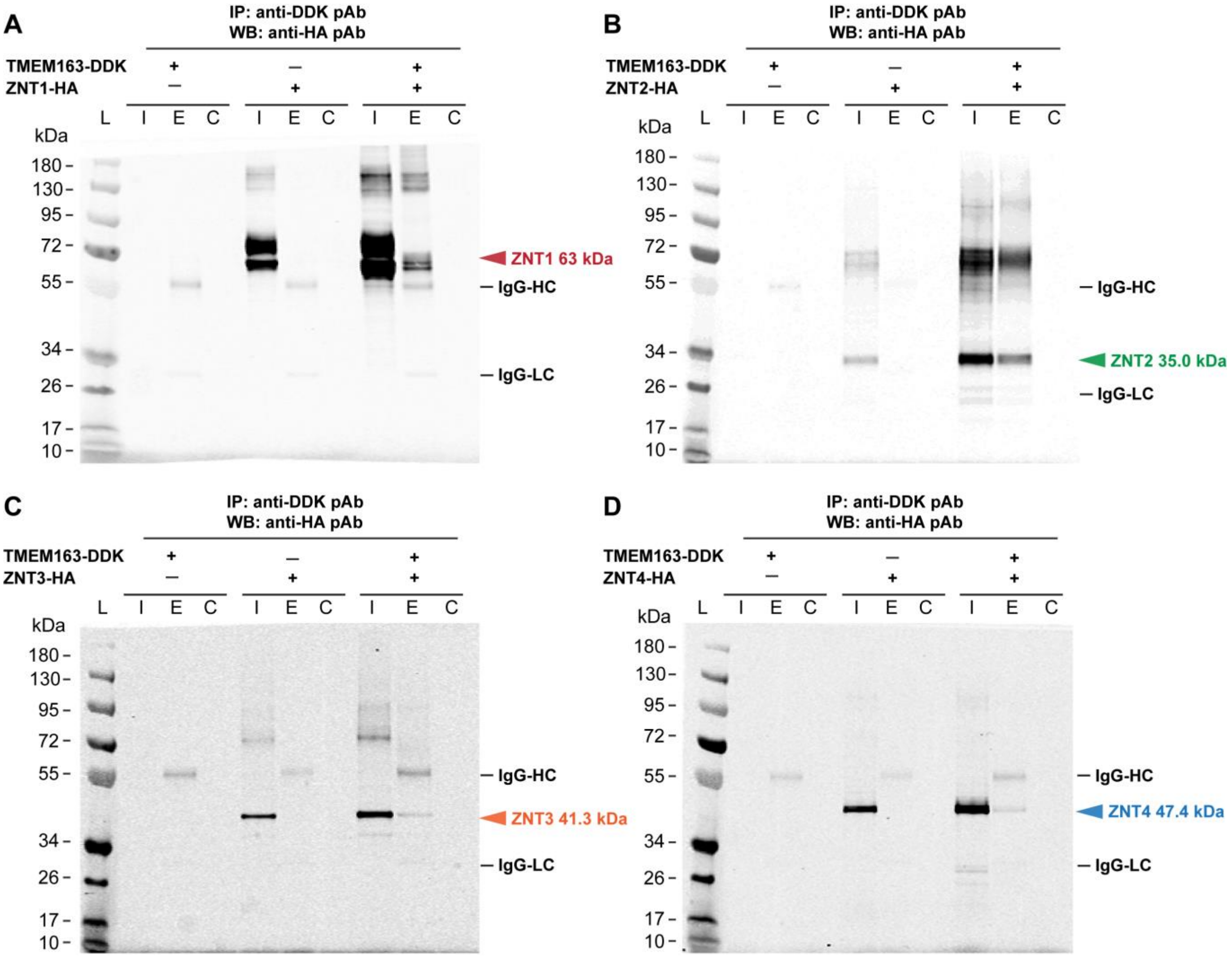


**Figure S2**. **Validations of co-immunoprecipitation experiments reveal specificity of protein-protein interaction.** To ensure that the proteins do not bind non-specifically to the agarose beads, all co-IP trials included control agarose resin. The resulting control lanes did not show discernible bands for all trials indicating specificity. Representative Western blot (WB) shows TMEM163-DDK co-immunoprecipitated (IP) with **A**) ZNT1-HA, **B**) ZNT2-HA, **C**) ZNT3-HA, and **D**) ZNT4-HA. Each respective ZNT protein co-elutes with TMEM163. Predicted molecular weight (MW) of human proteins: ZNT1 = 55.3-63.0 kDa (*arrowhead*); ZNT2 = 35.0 kDa (*arrowhead*); ZNT3 = 41.3 kDa (*arrowhead*); ZNT4 = 47.4 kDa (*arrowhead*); IgG-HC, immunoglobulin-G heavy chain (MW ≅ 55 kDa); IgG-LC, immunoglobulin-G light chain (MW ≅ 23 kDa). L, protein ladder; I, input/cell lysate; C, control resin; E, elution; mAb, monoclonal antibody; pAb, polyclonal antibody. The representative images come from N ≥ 3 independent trials.

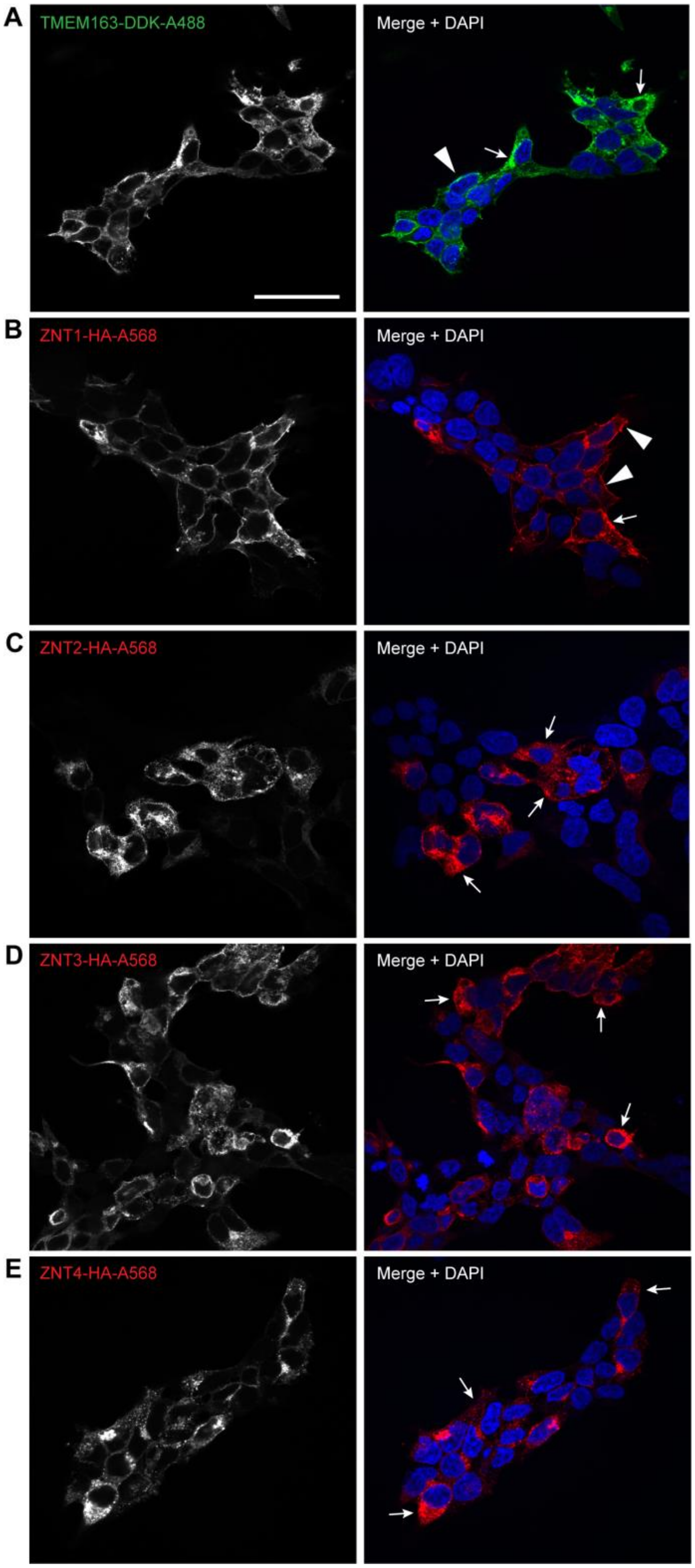


**Figure S3. Immunocytochemistry of heterologously expressed TMEM163 and ZNT proteins.** Representative confocal images of cells expressing **A**) TMEM163-DDK, **B**) ZNT1-HA, **C**) ZNT2-HA, **D**) ZNT3-HA, and **E**) ZNT4-HA. The DDK peptide tag was detected by anti-DDK mAb and visualized with Alexa-488 (green) anti-mouse secondary antibody. The HA peptide was detected by anti-HA pAb and visualized with Alexa-568 (red) anti-rabbit secondary antibody. DAPI stains the nuclei blue. TMEM163 and ZNT1 are detected within the plasma membrane (arrowhead) and intracellular compartments (arrow). Meanwhile, ZNT2, ZNT3, and ZNT4 proteins exhibit a punctate distribution pattern indicating localized in compartments (arrow). Scale bar: 100 $\mu$m.

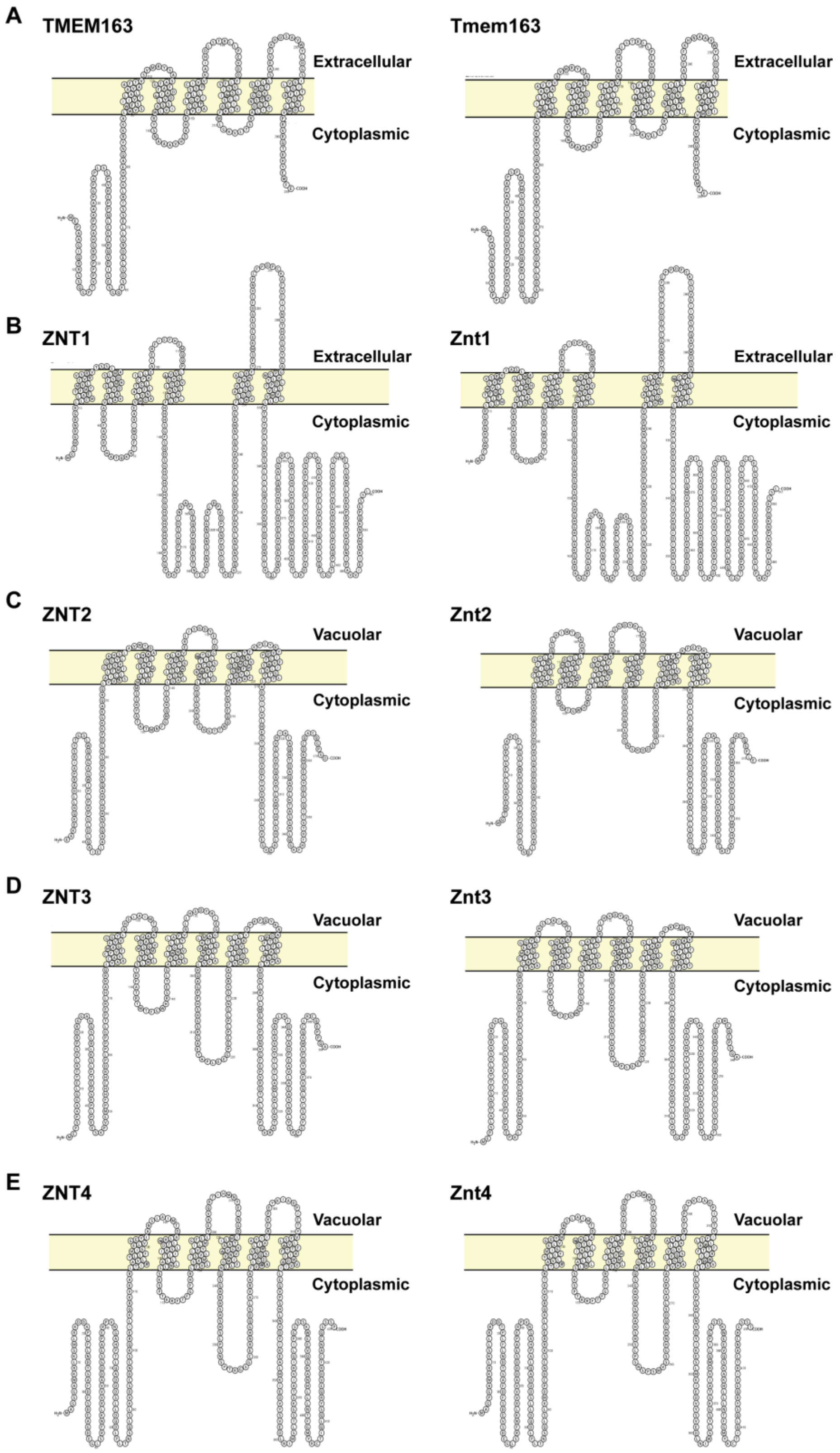


**Figure S4. Predicted protein secondary structures.** Human (left panel) and mouse (right panel) protein. **A**) TMEM163, **B**) ZNT1, **C**) ZNT2, **D**) ZNT3, and **E**) ZNT4. The diagrams were generated using the online Protter software (https://wlab.ethz.ch/protter/start/). Omasits U, Ahrens CH, Müller S, Wollscheid B. (2014). Bioinformatics, 30(6), 884-886. doi: 10.1093/bioinformatics/btt607.
